# Mice immunized with the vaccine candidate HexaPro spike produce neutralizing antibodies against SARS-CoV-2

**DOI:** 10.1101/2021.02.27.433054

**Authors:** Chotiwat Seephetdee, Nattawut Buasri, Kanit Bhukhai, Kitima Srisanga, Suwimon Manopwisedjaroen, Sarat Lertjintanakit, Nut Phueakphud, Chatbenja Pakiranay, Niwat Kangwanrangsan, Sirawat Srichatrapimuk, Somnuek Sungkanuparph, Suppachok Kirdlarp, Somchai Chutipongtanate, Arunee Thitithanyanont, Suradej Hongeng, Patompon Wongtrakoongate

## Abstract

Updated and revised versions of COVID-19 vaccines are vital due to genetic variations of the SARS-CoV-2 spike antigen. Furthermore, vaccines that are safe, cost-effective, and logistically friendly are critically needed for global equity, especially for middle to low income countries. Recombinant protein-based subunit vaccines against SARS-CoV-2 have been reported with the use of the receptor binding domain (RBD) and the prefusion spike trimers (S-2P). Recently, a new version of prefusion spike trimers, so called “HexaPro”, has been shown for its physical property to possess two RBD in the “up” conformation, as opposed to just one exposed RBD found in S-2P. Importantly, this HexaPro spike antigen is more stable than S-2P, raising its feasibility for global logistics and supply chain. Here, we report that the spike protein HexaPro offers a promising candidate for SARS-CoV-2 vaccine. Mice immunized by the recombinant HexaPro adjuvanted with aluminium hydroxide using a prime-boost regimen produced high-titer neutralizing antibodies for up to 56 days after initial immunization against live SARS-CoV-2 infection. In addition, the level of neutralization activity is comparable to that of convalescence sera. Our results indicate that the HexaPro subunit vaccine confers neutralization activity in sera collected from mice receiving the prime-boost regimen.

## Introduction

The coronavirus disease 2019 (COVID-19) caused by the novel coronavirus, severe acute respiratory syndrome coronavirus 2 (SARS-CoV-2), is a current global plaque. The incidence for this pandemic reported by World Health Organization (WHO) on January, 31, 2021 has included 102 million cumulative confirmed cases and over 2.2 million deaths globally. There is an urgent need for preventative vaccines and therapeutics. SARS-CoV-2 is an enveloped, single-stranded RNA virus. Its genome encodes four structural proteins comprising of spike (S), membrane glycoprotein (M), envelope (E), and nucleocapsid (N) proteins. The spike protein mediates viral entry by binding to the host receptor angiotensin-converting enzyme 2 (ACE2), via the receptor-binding domain (RBD). This interaction triggers a substantial conformational alteration of the spike from a prefusion conformation to a highly stable postfusion conformation [1–4]. The spike protein can induce production of neutralizing antibody in patients, indicating its immunogenic property. Thus, it has been widely adopted for vaccine development. However, the ongoing COVID-19 pandemic has led to SARS-CoV-2 spike variants with serious concerns such as D614G, N501Y, E484K, and 69/70 deletion [5–7]. Some of these variants can be highly transmissible and capable of escape vaccine-induced neutralizing antibody responses [8].

A key strategy for vaccine design against coronaviruses SARS-CoV and MERS-CoV has aimed at stabilizing the metastable prefusion conformation of the spike protein homologues [9,10]. The prefusion stabilization has been achieved by two consecutive proline substitutions (S-2P) in a turn between the central helix and heptad repeat 1 (HR1). These S-2P variants together with a C-terminus foldon trimerization domain have been shown as a superior immunogen [10]. As the consequence, the SARS-CoV-2 S-2P has been employed in currently used vaccines including mRNA-1273 [11], BNT162b2 [12], and ChAdOx1 [13].

In this research, we evaluated a potential COVID-19 subunit vaccine using a recently published prefusion-stabilized spike ectodomain namely HexaPro developed by McLellan and colleagues [14]. Here we show that the HexaPro subunit vaccine administered with aluminium hydroxide adjuvant in mice elicits strong neutralizing antibody response against SARS-CoV-2. This finding holds a promise towards a next-generation coronavirus vaccine development using the HexaPro spike protein.

## Results

### Expression and purification of recombinant SARS-CoV-2 HexaPro spike protein

The prefusion-stabilized HexaPro construct (Fig. 1A) encoding the spike ectodomain of SARS-CoV-2 with proline substitution at residues 817, 892, 899, 942, 986, and 987, “GSAS” substitution at residues 682-685 (the furin cleavage site), and C-terminal foldon trimerization motif [14] was used to produce the HexaPro subunit vaccine in HEK293 cells. Transient transfection of HexaPro-encoding plasmid into the cells resulted in expression of recombinant protein in culture supernatant. The recombinant HexaPro protein was purified by Ni-NTA chromatography followed by size exclusion chromatography. Purity of the purified recombinant HexaPro was ascertained by SDS-PAGE (Fig. 1B). Using pooled convalescence sera from COVID-19 patients and western blot analysis, we show that the HexaPro spike is immunogenic. To confirm the immunogenicity of the HexaPro spike, immunofluorescence staining was then performed and validated that the spike structural variant could be detected by antibodies against SARS-CoV-2 spike protein as well as by the pooled convalescent sera (Fig. 2). These results illustrate the potential of HexaPro recombinant protein as a subunit vaccine.

**Figure 1.**
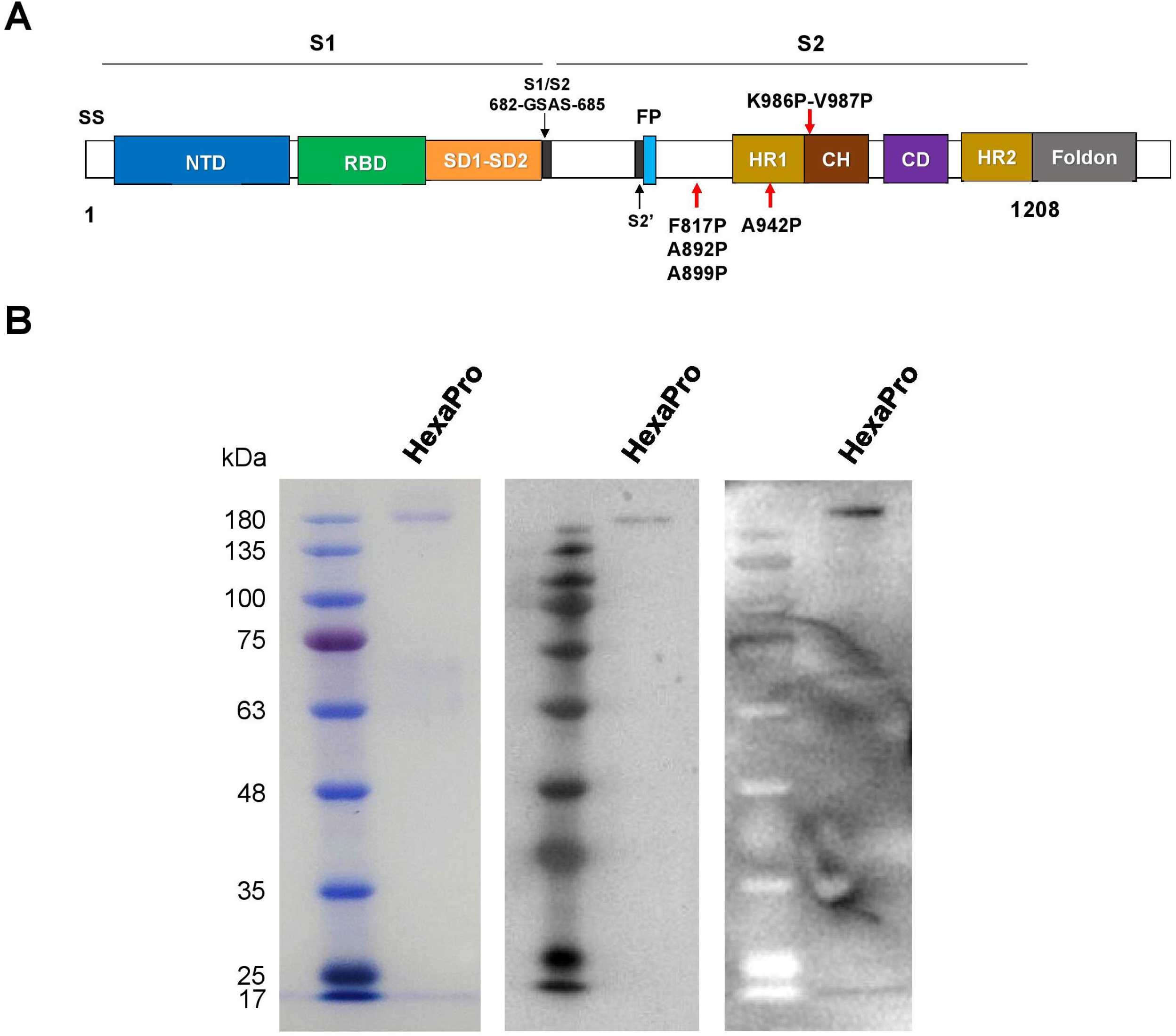
The recombinant SARS-CoV-2 HexaPro spike protein. (A) Schematic representation of the prefusion-stabilized SARS-CoV-2 HexaPro ectodomain showing the S1 and S2 subunits. Four additional proline substitutions from S-2P construct are indicated by the red arrows shown below the construct. (B) The HexaPro protein expressed in HEK293T cells was purified and characterized by SDS-PAGE (left), western blot using a commercial anti-RBD (middle), and western blot using pooled convalescence sera (right).

**Figure 2.**
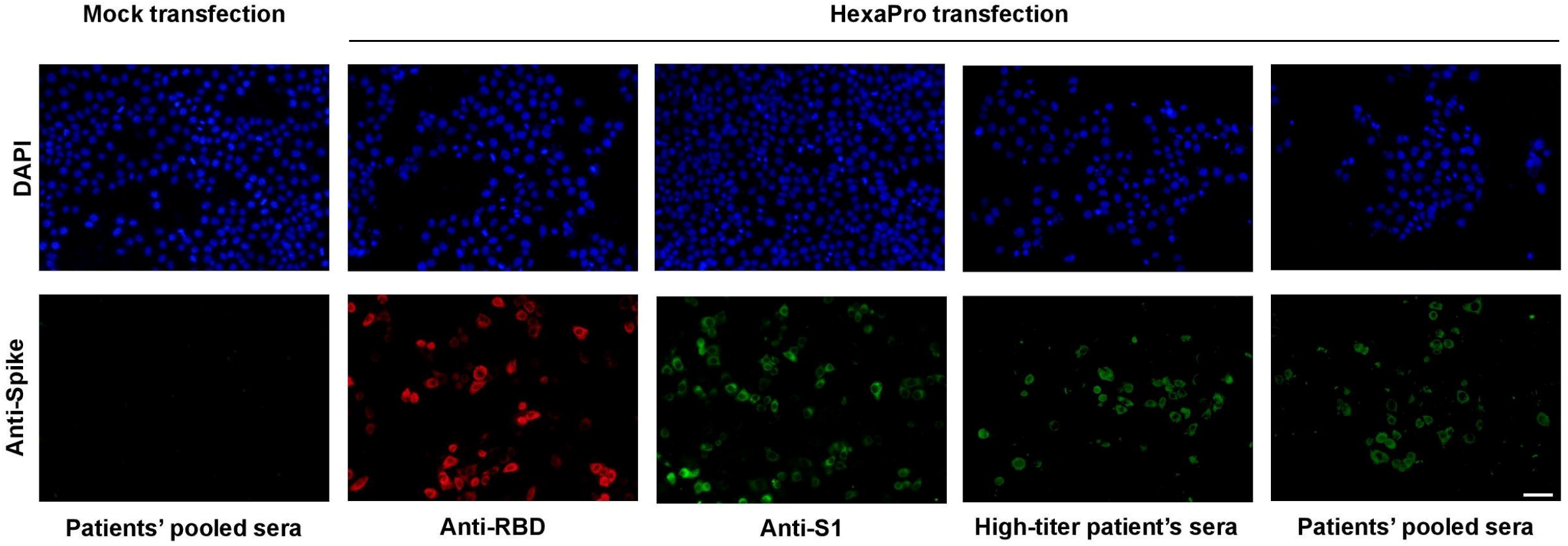
Immunofluorescence staining of the HexaPro spike expressed in HeLa cells using antibodies against spike RBD and S1 subunits, and convalescent sera derived from a patient with high-titer neutralization activity and from pooled sera. Scale bar, 50 µm.

### Neutralization of SARS-CoV-2 by sera collected from HexaPro-immunized mice

The HexaPro subunit vaccine was then evaluated for its immunization activity via neutralization of SARS-CoV-2 by immunized mouse sera. An immunization protocol of low priming dose followed by high booster dose was followed. At day 0, C57BL/C mice were prime-immunized with 1 µg of HexaPro adjuvanted with aluminium hydroxide (100 µg) via intramuscular administration. At day 21, the mice were boost-immunized with 5 µg of HexaPro (Fig. 3A). The microneutralization assay using live SARS-CoV-2 infection in Vero E6 cells was performed with sera collected at days 14, 35 and 56 after initial immunization. At 14 days after the priming dose, a minute neutralizing activity was observed in vaccinated mice (Fig. 3B). However, sera from vaccinated mice collected 14 days after the booster dose elicits high neutralization titers. Furthermore, the level of neutralization activity was sustained at least 56 days after the initial immunization (Fig. 3B). Together, our results indicate that the HexaPro subunit vaccine adjuvanted with aluminium hydroxide confers neutralization activity in sera collected from mice receiving the prime-boost regimen.

**Figure 3.**
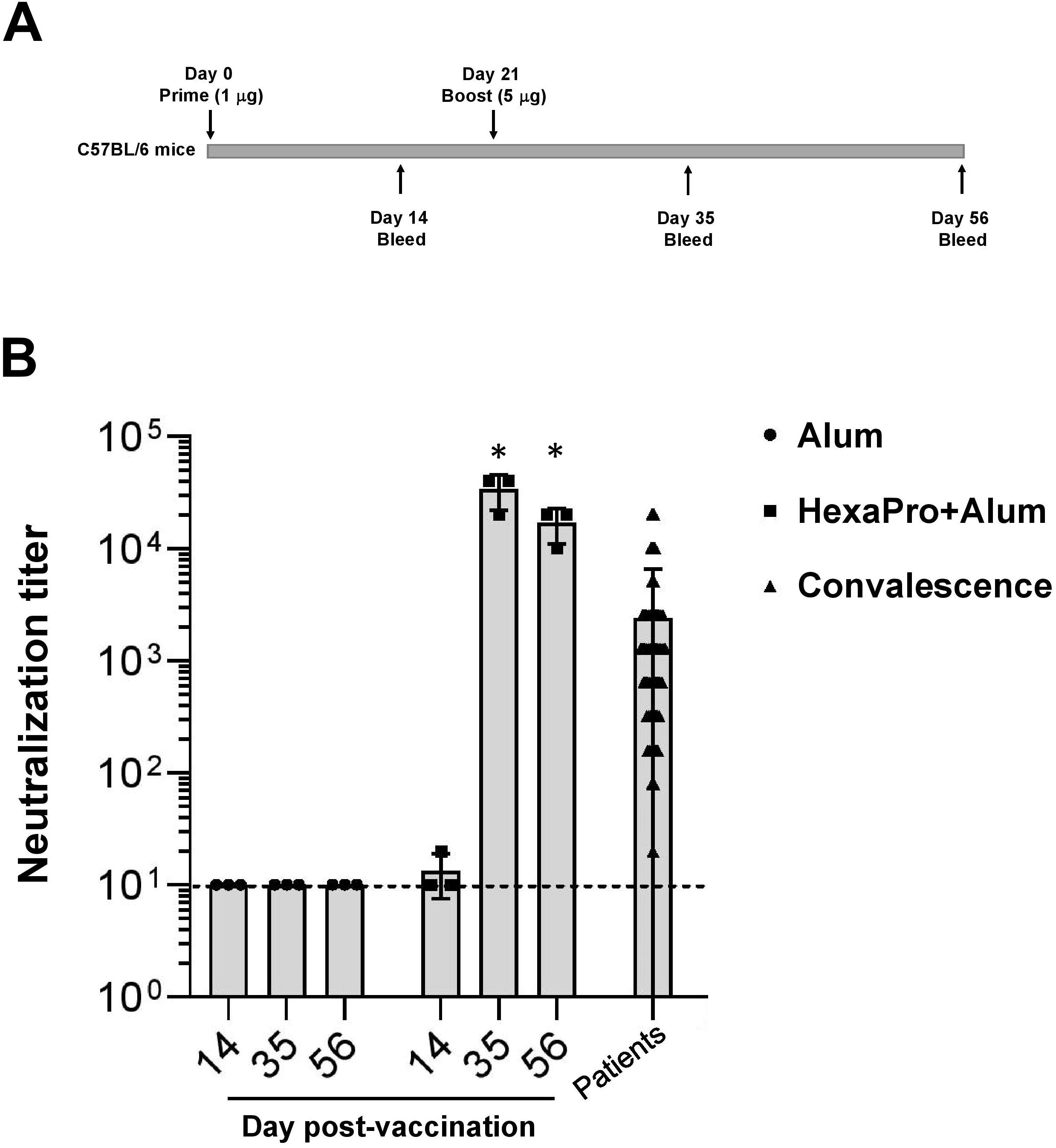
The prime-boost regimen using the recombinant HexaPro adjuvanted with aluminium hydroxide results in sera possessing neutralization activity. (A) C57BL/6 mice were vaccinated intramuscularly with Alum or HexaPro (1 µg)+Alum. At day 21 they then received a booster dose with HexaPro (5 µg)+Alum. (B) The virus neutralization endpoint titer of sera collected from mice and from convalescence sera. The dashed line shows the limit of detection. Neutralization activity at days 35 and 56 was compared to day 14. The error bars indicate the ±SD. Comparisons were performed by Student’s t-test (unpaired, two tail); *p < 0.01.

## Discussion

In this report, we observed a potent SARS-CoV-2 neutralizing activity delivered by the subunit vaccine HexaPro spike; four amino acids of which were substituted by McLellan and colleagues into beneficial prolines leading to a more stable spike variant [14]. Specifically, the amino acid substitution was engineered within the S2 domain of the original S-2P spike [2]. This novel prefusion variant possesses 30% of the spike trimers being an “up” conformation with two exposed RBD as opposed to just one exposed RBD found in S-2P. Due to its enhanced stability, the HexaPro spike has been proposed for its potential as a COVID-19 vaccine.

Using aluminium hydroxide in vaccination has been shown to enhance activation of inflammatory dendritic cells and T-cell responses [15–17]. In a phase 1 trial, alum was employed with the inactivated SARS-CoV-2 vaccine BBV152 [18] as well as in ongoing clinical trials of COVID-19 vaccines including subunit vaccines (NCT04522089; NCT04527575; NCT04683484; NCT04742738) and an inactivated SARS-CoV-2 vaccine (NCT0464148). In addition, we also adopted a regimen of prime-boost immunization using low priming dose followed by high booster dose. A number of studies and clinical trials have demonstrated that a lower priming dose, followed by a subsequent higher booster dose can induce greater levels of immune response [19–21] including the COVID-19 vaccine ChAdOx1 [13]. Importantly, effector cells are adversely induced by higher doses of antigen at prime immunization. On the other hand, immune memory cells are promisingly induced by lower doses at prime immunization making this regimen suitable for long-term immunological memory [22].

Altogether, due to its highly stable conformation, feasible production and logistic applicability, this HexaPro spike should be further developed into a COVID-19 vaccine, and exploited for its efficacy in viral challenge studies using different SARS-CoV-2 genetic variants. Moreover, mRNA, DNA, and viral vector vaccines should benefit by the use of this HexaPro variant.

## Materials and Methods

### Ethics statement

Mouse experiments were performed under the Animal Ethics approved by Faculty of Science, Mahidol University (MUSC63-016-524). PCR-confirmed COVID-19 patients (n=58) were hospitalized at Chakri Naruebodindra Medical Institute, Faculty of Medicine Ramathibodi Hospital, Mahidol University. Serum specimens were collected from patients 14 day-post infection. The study protocol was approved by Faculty of Medicine Ramathibodi Hospital.

### Expression and purification of HexaPro subunit vaccine

The mammalian expression plasmid containing SARS-CoV-2 HexaPro spike was obtained from Addgene (Addgene plasmid # 154754; http://n2t.net/addgene:154754; RRID: Addgene_154754). HEK293T cells were transiently transfected with the HexaPro plasmid by calcium phosphate transfection. Cells and culture medium were separated by centrifugation. Supernatant were concentrated with Amicon® Ultra–15 Ultrace–30K centrifugal filter unit (MERCK). Cell protein contents were extracted with a lysis buffer composed of 50 mm sodium phosphate, 300 mM NaCl, 20 mm imidazole, 1X cOmpleteTM EDTA-free protease inhibitor cocktail (Roche), 1 mM Phenylmethylsulfonyl fluoride (PMSF, Sigma-Aldrich), and 1% Triton-X (Sigma-Aldrich). Protein extracts were filtered through a 0.22 µm NalgeneTM syringe filter (Thermo ScientificTM). S HexaPro protein was then purified with HisTrap HP (cytiva) equilibrated with a buffer composed of 50 mm Sodium phosphate, 300 mM NaCl, and 20 mm imidazole. Fractions containing HexaPro were pooled and exchanged to phosphate-buffered saline (PBS). Purified protein was digested with HRV3C protease to remove purification tags. The protein was further purified with Sephacryl S-300 HR (GE Healthcare) with PBS. Fractions which contain HexaPro protein were pooled and analyzed with SDS-PAGE and western blot against the SARS-CoV-2 RBD protein (Sino Biological, Cat#40592-T62) or pooled convalescent sera. Purified proteins are kept in −80 °C until use.

### Immunofluorescence staining

HeLa cells were transiently transfected with the plasmid encoding HexaPro using lipofectamine 3000 (Invitrogen, Cat#L3000008). Cells were fixed with 4% PFA and were incubated with either a polyclonal antibody against the SARS-CoV-2 RBD protein (Sino Biological, Cat#40592-T62) or a monoclonal antibody against the SARS-CoV-2 S1 protein (MyBioSource, Cat#MBS434277). A goat anti-rabbit secondary antibody (IgG) conjugated with Alexa Fluor 594 (Invitrogen, Cat#A-11037) or a goat anti-mouse secondary antibody (IgG) conjugated with Alexa Fluor 488 (Invitrogen, Cat#A-11029) was used for visualization under a fluorescence microscope. For convalescent serum staining, cells were incubated with heat-inactivated serum and visualized with a goat anti-human secondary antibody (IgG) conjugated with FITC (Abcam, Cat#ab97224).

### Mouse immunization

Female C57BL/6 mice (7-9 weeks old, n = 3 per group) were ordered from Nomura Siam International. Mice were given a prime-boost immunization intramuscularly (i.m.) spaced three weeks apart. For antigen formulation, SARS-CoV-2 S HexaPro protein (1 µg for the first dose and 5 µg for the booster dose) was mixed with 100 µg of aluminium hydroxide (Invivogen, Cat#vac-alu-250). Serum was collected for analysis on study days 14, 35, and 56 after initial immunization.

### Microneutralization assay

Heat-inactivated sera at 56°C for 30 minutes were two-fold serially diluted, starting with a dilution of 1:10. The serum dilutions were mixed with equal volumes of 100 TCID50 of SARS-CoV-2. After 1 hr of incubation at 37°C, 100 μl of the virus–serum mixture at each dilution was added in duplicate to Vero E6 cell monolayers in a 96-well microtiter plate. The last two columns are set as virus control, cell control, and virus back-titration. The plates were incubated at 37°C in 5% CO_2_ in a humidified incubator. After 2 days of incubation, the medium was discarded and the cell monolayer was fixed with cold fixative (1:1 methanol:acetone) for 20 min on ice. Viral protein in virus-infected cell was detected by ELISA assay. The cells were washed 3 times with PBST before blocking with 2% BSA for 1 hr at room temperature. After washing, the viral nucleocapsid was detected using 1:5000 of SARS-CoV/SARS-CoV-2 Nucleocapsid monoclonal antibody (Sino Biological, Cat#40143-R001) by incubation at 37°C for 1 h. After removal of the detection antibody, 1:2000 HRP-conjugated goat anti-rabbit polyclonal antibody (Dako, Denmark A/S, Cat#P0448) was added and the plate was incubated at 37°C for 1 h. After washing, the TMB substrate (KPL, Cat#5120-0075) was added. After 10 min incubation, the reaction was stopped by the addition of 1N HCl. Optical density (O.D.) at 450 and 620 nm was measured by a microplate reader (Tecan Sunrise).

The virus neutralization endpoint titer of each serum was calculated using the following equation:

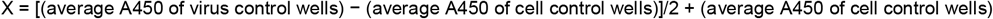

The reciprocal of the highest dilution of serum with O.D. values less than X is considered positive for neutralization activity. Serum samples tested negative at a dilution of 1:10 were assigned an NT titer of <10. The serum that is positive at 1:10 dilution will be reported as a NT titer of 20.

Each sample will be done in duplicate. All the experiments will be used live SARS-CoV–2 viruses at passage 3 or 4 with the Vero E6 cells at the maximum passages of 20. The activities with live viruses are carried in a certified biosafety level 3 facility.

## Acknowledgements

This research project was supported by Faculty of Medicine Ramathibodi Hospital, Mahidol University and CIF Grant, Faculty of Science, Mahidol University. CS, SL and CP were supported by Science Achievement Scholarship of Thailand. PW was supported by Mahidol University (New Discovery and Frontier Research Grant), and Office of National Higher Education Science Research and Innovation Policy Council by Program Management Unit for Human Resources and Institutional Development, Research and Innovation (PMU-B). Alum was kindly gifted by Dr. Waranyoo Phoolcharoen. We are indebted to COVID-19 Cluster and PW lab members for their suggestions and comments.

## Notes

### Competing Interest Statement

The authors have declared no competing interest.

## References

1. Hoffmann, M.; Kleine-Weber, H.; Schroeder, S.; Kruger, N.; Herrler, T.; Erichsen, S.; Schiergens, T.S.; Herrler, G.; Wu, N.H.; Nitsche, A., et al. SARS-CoV-2 Cell Entry Depends on ACE2 and TMPRSS2 and Is Blocked by a Clinically Proven Protease Inhibitor. Cell 2020, 10.1016/j.cell.2020.02.052, doi:10.1016/j.cell.2020.02.052

2. Wrapp, D.; Wang, N.; Corbett, K.S.; Goldsmith, J.A.; Hsieh, C.L.; Abiona, O.; Graham, B.S.; McLellan, J.S. Cryo-EM structure of the 2019-nCoV spike in the prefusion conformation. Science (New York, N.Y.) 2020, 367, 1260–1263.

3. Yan, R.; Zhang, Y.; Li, Y.; Xia, L.; Guo, Y.; Zhou, Q. Structural basis for the recognition of SARS-CoV-2 by full-length human ACE2. Science (New York, N.Y.) 2020, 367, 1444–1448.

4. Walls, A.C.; Park, Y.J.; Tortorici, M.A.; Wall, A.; McGuire, A.T.; Veesler, D. Structure, Function, and Antigenicity of the SARS-CoV-2 Spike Glycoprotein. Cell 2020, 181, 281–292 e286.

5. Ozono, S.; Zhang, Y.; Ode, H.; Sano, K.; Tan, T.S.; Imai, K.; Miyoshi, K.; Kishigami, S.; Ueno, T.; Iwatani, Y., et al. SARS-CoV-2 D614G spike mutation increases entry efficiency with enhanced ACE2-binding affinity. Nat Commun 2021, 12, 848.

6. Wang, P.; Liu, L.; Iketani, S.; Luo, Y.; Guo, Y.; Wang, M.; Yu, J.; Zhang, B.; Kwong, P.D.; Graham, B.S., et al. Increased Resistance of SARS-CoV-2 Variants B.1.351 and B.1.1.7 to Antibody Neutralization. bioRxiv 2021.

7. Liu, Z.; VanBlargan, L.A.; Bloyet, L.M.; Rothlauf, P.W.; Chen, R.E.; Stumpf, S.; Zhao, H.; Errico, J.M.; Theel, E.S.; Liebeskind, M.J., et al. Identification of SARS-CoV-2 spike mutations that attenuate monoclonal and serum antibody neutralization. Cell host & microbe 2021.

8. Wang, Z.; Schmidt, F.; Weisblum, Y.; Muecksch, F.; Barnes, C.O.; Finkin, S.; Schaefer-Babajew, D.; Cipolla, M.; Gaebler, C.; Lieberman, J.A., et al. mRNA vaccine-elicited antibodies to SARS-CoV-2 and circulating variants. Nature 2021, 10.1038/s41586-021-03324-6, doi:10.1038/s41586-021-03324-6

9. Kirchdoerfer, R.N.; Cottrell, C.A.; Wang, N.; Pallesen, J.; Yassine, H.M.; Turner, H.L.; Corbett, K.S.; Graham, B.S.; McLellan, J.S.; Ward, A.B. Pre-fusion structure of a human coronavirus spike protein. Nature 2016, 531, 118–121.

10. Pallesen, J.; Wang, N.; Corbett, K.S.; Wrapp, D.; Kirchdoerfer, R.N.; Turner, H.L.; Cottrell, C.A.; Becker, M.M.; Wang, L.; Shi, W., et al. Immunogenicity and structures of a rationally designed prefusion MERS-CoV spike antigen. Proc Natl Acad Sci U S A 2017, 114, E7348–E7357.

11. Baden, L.R.; El Sahly, H.M.; Essink, B.; Kotloff, K.; Frey, S.; Novak, R.; Diemert, D.; Spector, S.A.; Rouphael, N.; Creech, C.B., et al. Efficacy and Safety of the mRNA-1273 SARS-CoV-2 Vaccine. The New England journal of medicine 2021, 384, 403–416.

12. Polack, F.P.; Thomas, S.J.; Kitchin, N.; Absalon, J.; Gurtman, A.; Lockhart, S.; Perez, J.L.; Pérez Marc, G.; Moreira, E.D.; Zerbini, C., et al. Safety and Efficacy of the BNT162b2 mRNA Covid-19 Vaccine. The New England journal of medicine 2020, 383, 2603–2615.

13. Voysey, M.; Clemens, S.A.C.; Madhi, S.A.; Weckx, L.Y.; Folegatti, P.M.; Aley, P.K.; Angus, B.; Baillie, V.L.; Barnabas, S.L.; Bhorat, Q.E., et al. Safety and efficacy of the ChAdOx1 nCoV-19 vaccine (AZD1222) against SARS-CoV-2: an interim analysis of four randomised controlled trials in Brazil, South Africa, and the UK. Lancet (London, England) 2021, 397, 99–111.

14. Hsieh, C.L.; Goldsmith, J.A.; Schaub, J.M.; DiVenere, A.M.; Kuo, H.C.; Javanmardi, K.; Le, K.C.; Wrapp, D.; Lee, A.G.; Liu, Y., et al. Structure-based design of prefusion-stabilized SARS-CoV-2 spikes. Science (New York, N.Y.) 2020, 369, 1501–1505.

15. Kool, M.; Soullié, T.; van Nimwegen, M.; Willart, M.A.; Muskens, F.; Jung, S.; Hoogsteden, H.C.; Hammad, H.; Lambrecht, B.N. Alum adjuvant boosts adaptive immunity by inducing uric acid and activating inflammatory dendritic cells. J Exp Med 2008, 205, 869–882.

16. Kool, M.; Pétrilli, V.; De Smedt, T.; Rolaz, A.; Hammad, H.; van Nimwegen, M.; Bergen, I.M.; Castillo, R.; Lambrecht, B.N.; Tschopp, J. Cutting edge: alum adjuvant stimulates inflammatory dendritic cells through activation of the NALP3 inflammasome. Journal of immunology (Baltimore, Md.: 1950) 2008, 181, 3755–3759, doi:10.4049/jimmunol.181.6.3755

17. HogenEsch, H.; O’Hagan, D.T.; Fox, C.B. Optimizing the utilization of aluminum adjuvants in vaccines: you might just get what you want. NPJ Vaccines 2018, 3, 51.

18. Ella, R.; Vadrevu, K.M.; Jogdand, H.; Prasad, S.; Reddy, S.; Sarangi, V.; Ganneru, B.; Sapkal, G.; Yadav, P.; Abraham, P., et al. Safety and immunogenicity of an inactivated SARS-CoV-2 vaccine, BBV152: a double-blind, randomised, phase 1 trial. Lancet Infect Dis 2021.

19. Pace, D.; Khatami, A.; McKenna, J.; Campbell, D.; Attard-Montalto, S.; Birks, J.; Voysey, M.; White, C.; Finn, A.; Macloed, E., et al. Immunogenicity of reduced dose priming schedules of serogroup C meningococcal conjugate vaccine followed by booster at 12 months in infants: open label randomised controlled trial. BMJ 2015, 350, h1554.

20. Nitayaphan, S.; Pitisuttithum, P.; Karnasuta, C.; Eamsila, C.; de Souza, M.; Morgan, P.; Polonis, V.; Benenson, M.; VanCott, T.; Ratto-Kim, S., et al. Safety and immunogenicity of an HIV subtype B and E prime-boost vaccine combination in HIV-negative Thai adults. J Infect Dis 2004, 190, 702–706, doi:10.1086/422258

21. Sandström, E.; Nilsson, C.; Hejdeman, B.; Bråve, A.; Bratt, G.; Robb, M.; Cox, J.; Vancott, T.; Marovich, M.; Stout, R., et al. Broad immunogenicity of a multigene, multiclade HIV-1 DNA vaccine boosted with heterologous HIV-1 recombinant modified vaccinia virus Ankara. J Infect Dis 2008, 198, 1482–1490.

22. Kardani, K.; Bolhassani, A.; Shahbazi, S. Prime-boost vaccine strategy against viral infections: Mechanisms and benefits. Vaccine 2016, 34, 413–423, doi:10.1016/j.vaccine.2015.11.062

